# No evidence for cross-contextual consistency in spatial learning and behavioural flexibility in a passerine

**DOI:** 10.1101/2020.09.04.282566

**Authors:** CA Troisi, AC Cooke, GL Davidson, I de la Hera, MS Reichert, JL Quinn

**Affiliations:** School of Biology, Earth and Environmental Sciences, University College Cork, Ireland; Department of Psychology Department, University of Cambridge, UK; Department of Integrative Biology, Oklahoma State University, USA

**Keywords:** cognition, spatial learning, inhibitory control, great tits, consistency, repeatability

## Abstract

Although the evolution of cognitive differences among species has long been of interest in ecology, whether natural selection acts on cognitive processes within populations has only begun to receive similar attention. One of the key challenges is to understand how consistently cognitive traits within any one domain are expressed over time and across different contexts, as this has direct implications for the way in which selection might act on this variation. Animal studies typically measure a cognitive domain using only one task in one context, and assume that this captures the likely expression of that domain in different contexts. This deficit is not surprising because, from an ecologist’s perspective, cognitive tasks are notoriously laborious to employ, and for design reasons most tasks can only be deployed in a specific context. Thus our knowledge of whether individual differences in cognitive abilities are consistent across contexts is limited. Using a wild population of great tits (*Parus major*) we characterised consistency of two cognitive abilities, each in two different contexts: 1) spatial learning at two different spatial scales, and 2) behavioural flexibility as both performance in a detour reaching task and reversal learning in a spatial task. We found no evidence of a correlation between the two spatial learning speeds, or between the two measures of behavioural flexibility. This suggests that cognitive performance is highly plastic and sensitive to differences across tasks, or that variants of these well-known tasks may tap into different combinations of both cognitive and non-cognitive mechanisms, or that they simply do not adequately measure each putative cognitive domain. Our results highlight the challenges of developing standardised cognitive assays to explain natural behaviour and to understand the selective consequences of that variation.

## Introduction

The importance of evolutionary processes in explaining why individuals vary in their cognitive abilities is an emerging question in behavioural ecology. On the one hand, support for natural selection is accumulating in studies that show links between correlates of fitness and cognitive measures (Ashton et al., 2018; Cauchard et al., 2013; Cole et al., 2012; Cole & Quinn, 2012; Keagy et al., 2009; Raine & Chittka, 2008). On the other, it remains unclear whether selection on individual variation in cognitive traits will result in a meaningful response (Shaw & Schmelz, 2017). One reason for this uncertainty over the evolutionary consequences of selection on cognition is that estimates of heritability are almost entirely lacking from natural populations (Langley et al., 2020; Quinn et al., 2016; reviewed in Croston et al., 2015), partly because generating reliable estimates of heritability demand large pedigrees (Quinn et al., 2006), and instead most researchers are forced to accept the phenotypic gambit, i.e., to assume that if a trait is repeatable, it is likely to be heritable, or that if two traits are phenotypically correlated, they are also genetically correlated (but see Quinn et al., 2016).

A recent meta-analysis has shown that repeatability among cognitive traits is highly variable, and that behaviours in certain types of cognitive tasks are less repeatable than others, thus providing less reliable measures of cognition (Cauchoix et al., 2018). This is particularly the case for contextual repeatability whereby two different tasks or the same task in different contexts aim to test the same cognitive trait, as opposed to temporal repeatability in which the exact same task is repeated over time (Cauchoix et al., 2018). Robust measures of individual differences in cognitive abilities are generally lacking, particularly within the same cognitive domain (i.e. statistically derived group of factors that capture the variance in a set of tasks) (Shaw & Schmelz, 2017; van Horik, Langley, Whiteside, Laker, & Madden, 2018). In fact, other studies looking at the same domain-specific cognitive abilities found little evidence of contextual repeatability, particularly for associative learning (Boogert et al., 2011; Bray et al., 2014; Brucks et al., 2017; Guenther & Brust, 2017; Isden et al., 2013; Keagy et al., 2009; Morand-Ferron et al., 2011; Shaw et al., 2015; van Horik et al., 2019; van Horik, Langley, Whiteside, Laker, Beardsworth, et al., 2018; van Horik, Langley, Whiteside, Laker, & Madden, 2018; Vernouillet et al., 2018). While a handful of studies have adopted test batteries to measure individual differences in cognitive abilities (Anderson et al., 2017; Ashton et al., 2018; Boogert et al., 2011; Guenther & Brust, 2017; Isden et al., 2013; Shaw et al., 2015; Soha et al., 2019; van Horik, Langley, Whiteside, Laker, & Madden, 2018), this approach remains relatively rare and few studies have explicitly tested contextual repeatability in multiple cognitive abilities. The gold standard would be to measure multiple traits across time and contexts in order to validate those measures of cognition (Völter et al., 2018). In this study we examined performance of two cognitive traits of key functional significance in behavioural ecology, spatial learning across contexts (i.e. spatial scale) and behavioural flexibility across two types of cognitive tasks.

Spatial learning, a form of learning relating to information about orientation and location, is a fundamental cognitive process that affects many aspects of an animal’s ecology. For example, spatial learning helps individuals to find food and mates, to monitor their territory boundaries, and at a larger scale to navigate their migration routes (Healy & Hurly, 2004). Several processes are involved in different navigational methods or attention to different cues. Animals can use view-matching strategies, such as landmark-matching or panorama-matching, where the apparent size or distance of landmarks, or the shape of the surroundings are important in matching a remembered view (Pritchard et al., 2018; Pritchard & Healy, 2018). Animals may also use strategies where the absolute distance or direction of a goal from a landmark is of importance (Pritchard et al., 2018; Pritchard & Healy, 2018). Moreover, these methods are not mutually exclusive and can be applied simultaneously in different environments or during different stages of navigation (Pritchard & Healy, 2018). In fact, evidence in the field and in captivity suggests that individuals rely on different information depending on the scale of their environment or the size of their enclosure (Chiandetti et al., 2007; Healy & Hurly, 1998; Sovrano et al., 2005, 2006). Although spatial learning is commonly referred to as a domain specific cognitive trait, such results question whether spatial learning should in fact be broken down into even more specific mechanisms. In humans, spatial abilities measured at different scales, have been shown to be underlined by some common processes for encoding, maintaining and transforming spatial representation, as well as some unique processes not shared at different scales of space (Hegarty et al., 2006; Montello, 1993). Yet the majority of studies in non-human animals adopt tasks that are relatively small in scale and do not differ in context (e.g. Branch et al., 2019; Sewall et al., 2013; Sonnenberg et al., 2019). If animals use different cues depending on the spatial scale, we may expect performance in a spatial learning task to differ across contexts. Nevertheless, the use of different navigational mechanisms does not preclude individual consistency across contexts, which would be suggestive of a meaningful trait that has the potential to be heritable.

Another cognitive trait that has received a lot of attention in cognitive ecology is behavioural flexibility, which allows individuals to adapt their behaviour to changes in their environment (Brown & Tait, 2014). In psychology, behavioural flexibility refers specifically to attentional shifting, rule switching and response reversal (Brown & Tait, 2014). Recently, behavioural and cognitive ecologists have been criticised for adopting the term to broadly describe any flexible behaviour, thus grouping behaviours that may be guided by different cognitive mechanisms (Audet & Lefebvre, 2017). As such, different assays of behavioural flexibility may involve different cognitive mechanisms, therefore limiting our ability to make broad inferences about what an animal’s performance in a particular context might mean for adaptive responses in the wild. Two tasks have been frequently used to test how well animals respond to changes in their environment – the detour reaching and reverse learning tasks. The detour task is thought to measure inhibitory control - an executive cognitive function that determines the ability to overcome a prepotent but disadvantageous response in favour of a more advantageous but less instinctive response. In this task animals must avoid a transparent barrier by inhibiting a motor response to go the most direct route towards a reward and instead move around the barrier (Boogert et al., 2011; MacLean et al., 2014). Reversal learning tasks are thought to measure how flexibly animals can adjust to changes in learned contingencies, whereby a novel response is rewarded and the previously rewarded response is not. While reverse learning can involve inhibitory control (Bari & Robbins, 2013) (particularly on the first reversal as opposed to multiple reversals in which rules are formed), it also involves instrumental conditioning (extinction and relearning of reinforced stimuli) (Brown & Tait, 2014). Previous work has directly tested the relationship between individual performance in a detour reach and a reversal learning task, with mixed results (Anderson et al., 2017; Ashton et al., 2018; Boogert et al., 2011; Brucks et al., 2017; Shaw et al., 2015; Soha et al., 2019; van Horik, Langley, Whiteside, Laker, & Madden, 2018). Moreover, all of the reversal learning tasks were based on object or colour associations, so it is still unclear how detour reach performance relates to reversal learning in a spatial context. Therefore, there is good reason to expect overlap in a domain general cognitive mechanism (i.e. inhibitory control) for both detour reaching and reverse learning, but there are also potentially some mechanisms that are exclusive to one task only, raising uncertainty as to whether one might expect these two traits to predict similar behaviours.

The great tit (*Parus major*) is a model species for ecological and behavioural studies (e.g. Aplin et al., 2015; Cole et al., 2011; Dutour et al., 2020; Loukola et al., 2020; Morand-Ferron et al., 2011). Great tits adapt well to temporary captivity, allowing for their use in controlled experiments on individual differences. Here we investigated consistency in spatial learning and behavioural flexibility across contexts. We measured spatial learning at two different spatial scales: at a large scale within an experimental room, and at a smaller scale within the birds’ individual home cages. We also measured behavioural flexibility across two different tasks (also at different scales): reversal learning, using a spatial feeder array in the experimental room and a detour apparatus in the home cage (MacLean et al., 2014). Our study had two objectives: first, to examine whether a learning task conducted in a home cage predicts measures of the same putative cognitive domain at a larger scale. Second, to investigate whether behavioural flexibility measured by the detour reach task predicts the behaviour of birds in a reversal learning task. If learning at different spatial scales is underlined by a general cognitive mechanism, then we would expect our measure of spatial learning in the home cage to be correlated to our measure of spatial learning in the experimental room. By contrast, if spatial learning at different scales requires different mechanisms, or attention to different cues, then we would not expect such a correlation. Similarly, if behavioural flexibility is underlined by inhibitory control, then we would expect measures coming from our two tasks to be correlated. However, if behavioural flexibility is a consolidated measure of different cognitive processes, then we expected measures emerging from those tasks be uncorrelated with each other (Miyake et al., 2000).

## Methods

Wild-caught great tits (n= 36) from the Bandon Valley, County Cork, Ireland, were brought into captivity, and later released at their capture site upon finishing the experiments (O’Shea et al., 2018). Birds were fitted with BTO rings for individual identification and a Passive Integrated Transponder (PIT) tag. Birds were housed in 57(h) × 56(d) × 46(w) cm plywood cages with two perches each and with an internal light set from 7:30 to 18:00. Birds had *ad libitum* access to food and water. Food consisted of sunflower hearts, peanuts, mealworms and waxworms. Out of the 36 birds brought into captivity, 28 birds learned the large-scale spatial learning task, and 25 of those also learned the reverse learning task. Out of the 36 birds, 30 learned the small-scale spatial learning task, and 32 completed all 10 trials of the detour reach task. Data collection took place between January and March 2019, and the mean time that birds were in captivity was 12 days.

### Large-scale spatial learning task

The large-scale spatial learning task took place in an experimental room of 460 (W) × 310 (L) × 265 (H) cm, with four feeders containing sunflower seeds placed in a square with sides of one meter, and a small plastic Christmas tree (150 cm high) placed in the centre for birds to rest and hide. Feeders were equipped with RFID readers to remotely log each visit by detecting the individual’s PIT tag. The task consisted of a habituation phase, training and test phases. Each trial within a phase lasted 1 hour, and birds were food deprived 1 hour beforehand. Each phase took place at least 12 hours apart (i.e. typically the following day). In the habituation phase, birds were released into the experimental room from a small opening in their home cage. Food was accessible and visible in all four feeders. Birds had to eat 10 seeds within 1 hour (based on seed husk collected on the floor after the trial) before progressing to the training phase. In the training phase, birds were once again released into the experimental room, but this time the seeds in each feeder were concealed by an opaque paper sheet so the food was no longer visible by the birds except from the RFID reader platform, from which it was also accessible. Once a bird visited any of the four feeders a total of 10 times (based on logged visits from RFID reader), they were advanced to the initial learning phase of the testing trials. During the testing trials, all feeders remained wrapped in paper but only one (randomly assigned) feeder contained food. The criterion for having learned the feeder position was to have visited the correct feeder 8 times within a moving window of 10 visits (Guenther & Brust, 2017). The number of visits to reach criterion was used as a measure of learning. Once the birds met the criterion, they were advanced to the reverse learning phase, in which a new feeder was allocated as the rewarded feeder. The same criterion was used as for the initial learning phase: 8 visits to the correct feeder within a moving window of 10 visits.

In all phases, birds were given one hour to visit the feeders before being returned to their home cage. Data from the loggers were reviewed on the same day to determine whether birds had met the criterion. If the birds did not reach their criterion within 6 trials, they were excluded as non-learners.

These data were collected as part of another experiment (Cooke et al., In prep) in which birds were exposed to different levels of simulated predation risk (treatment), during both the initial learning and reversal learning part of the experiment. We found no evidence that treatment in the previous experiment (Cooke et al., In prep) affected behaviour in the current one, so we analysed all individuals together (but see supplementary material for analysis of birds with no perceived predation risk (control birds)).

### Small-scale spatial learning task

Individuals were given artificial food items designed to mimic seeds/insect prey enclosed in an outer shell (Ihalainen et al., 2007). In our study, these artificial food items consisted of a sunflower seed encased in a paper parcel (1.8 cm × 1.8 cm). Adapting from Ihalainen et al. (2007), all birds were trained to handle the artificial food in their home cages in four steps in which they had to consume the seeds before advancing to the next step: (i) five food items with the seed sticking out from each parcel; (ii) five food items with the seed inside each parcel, but with a hole in the middle showing the seed inside; (iii) five food items with the seed completely hidden inside each parcel; and finally (iv) five food items with the seed completely hidden inside of each parcel with three rewarded (i.e. seed) and two unrewarded (i.e. made the reward inaccessible once the parcel was opened by wrapping the seed in duct tape). In this last step shielded items were used so that the birds would learn that not all parcels had accessible food. The birds had to eat all items before the training progressed to the next phase, or eat three rewarded food items and open two unrewarded food items in the last phase of training to proceed to the testing phase. In each training step parcels were placed centrally in the home cage in a small dish.

The testing phase consisted of ten parcels placed in each corner of the cage, in a small dish. Three of those locations contained only parcels that were unrewarded, while all parcels from the rewarded location contained seeds. The rewarded corner was allocated to each bird randomly. Every time the bird made 10 choices, irrespective of their location, each corner was rebaited to have 10 total parcels, so that the amount of parcels at each corner would not act as a cue for the bird.

During training and testing, birds had no *ad libitum* access to food, and only had access to food through the parcels. Individuals were not food deprived beforehand to allow for longer training and testing sessions. Water was still available *ad libitum*. Birds were trained and tested for a maximum of three hours consecutively. After three hours their *ad libitum* food was replaced in their cages. During training and testing, if a bird had not eaten any food for the last 1.5 hours, or less than 10 seeds for the last 2 hours, then the training/testing was stopped, and *ad libitum* food was placed back in their cages.

Trials stopped either once the bird had learned where to find seeds, based on a criterion of 8 correct visits in a moving window of 10 visits, or if they were not eating enough (see above). The number of choices to reach criterion was used as a measure of learning.

### Detour reach task

For this task, birds were required to retrieve a waxworm from inside a transparent cylinder, requiring them to make a detour around the cylinder to obtain the reward (Boogert et al., 2011; MacLean et al., 2014). The cylinder (3 cm length, 3.5 cm diameter) was made from plastic sheeting, open at both ends, and glued onto a cardboard base (7 cm × 20 cm). We also added a small perch (8 cm wide, 8 cm high) parallel to the cylinder, to avoid any biases in approach direction. The task had three phases: habituation, training and testing, and a waxworm was used as a reward in all phases. During the habituation and training phases the cylinder was opaque (black plastic), whereas it was transparent during the test. Before habituation and training birds were food deprived for 1 hour. For habituation, training and testing birds had *ad libitum* access to water and only had access to food through the task.

Birds were first habituated to the apparatus to reduce their fear towards the novel apparatus. For habituation birds were required to eat a waxworm placed in front of the opaque cylinder. Once they had completed this three times, birds advanced to the training phase of the experiment. During training birds were required to eat a waxworm placed in the middle of the opaque apparatus, by reaching around the cylinder into the open end without touching the exterior of the tube. Birds had to complete this four times to advance to the next phase. During the test phase, birds were presented the transparent apparatus with a waxworm placed in the middle. Birds were scored either a success (obtaining the worm without touching the tube) or a failure (touching the tube) and the tube was removed from the cage. This was repeated ten times in succession. Performance on this task was quantified as the number of successes out of ten. The test phase always occurred at least the day after the training phase was completed.

### Statistical Analysis

All analyses were conducted in R (R Core Team, 2019). Data followed a Poisson distribution, and therefore Kendall’s Tau correlations were conducted for non-normal distributions. We conducted a correlation test between the learning speed at the large and small scale, and between the performance on the detour reach task and reversal learning. In all analyses we excluded birds that learned on their first choice, as we cannot say whether they learned or made a choice by chance (large-scale spatial learning task n = 1; large-scale spatial reversal learning task n = 3; small-scale spatial learning task n =2) (but see supplementary material for analysis of all birds). For the two spatial learning tasks, we also compared overall learning speeds in terms of number of visits, and time, using generalised linear mixed-models with a Poisson error distribution and log function (Bates et al., 2015). Model assumptions were checked using DHARMa (Hartig, 2020).

### Data Availability

The dataset analysed during the current study, and the R code used to analyse them are available as supplementary material, and will be made available on the Open Science Framework upon publication.

### Ethics

We performed the experiment in accordance with the Association for the Study of Animal Behaviour guidelines, and the Animal Experimentation Ethics Committee of the University College Cork approved the study, under the number 2014/014 “The evolutionary and behavioural ecology of birds”. The Health Products Regulatory Authority approved the ethics for the project number AE19130/P017, and the National Park and Wildlife Services approved the capture of birds under the licence C01/2019.

## Results

On average, birds made 30.07 (SE = 4.48) choices before learning the large-scale spatial learning task (N=27), and 26.04 (SE = 3.09) choices before learning the small-scale spatial learning task (N=28). Although birds needed to make more choices to learn the large-scale spatial learning task (model estimate = 0.27; C.I. = 0.15-0.37; p-value <0.001), they took slightly less time (seconds) to learn it (model estimate = −0.03; C.I. =−0.03; −0.02; p-value <0.001), compared to the small-scale spatial learning task. Birds took on average 131.98 (SE = 12.29) minutes to learn the large-scale spatial learning task, while it took them 139.64 (SE = 19.91) minutes to learn the small-scale spatial learning task. There was no evidence of a correlation between the two measures (z = 0.22; tau = 0.03; p = 0.823, n=24; Figure 1.A).

**Figure 1:**
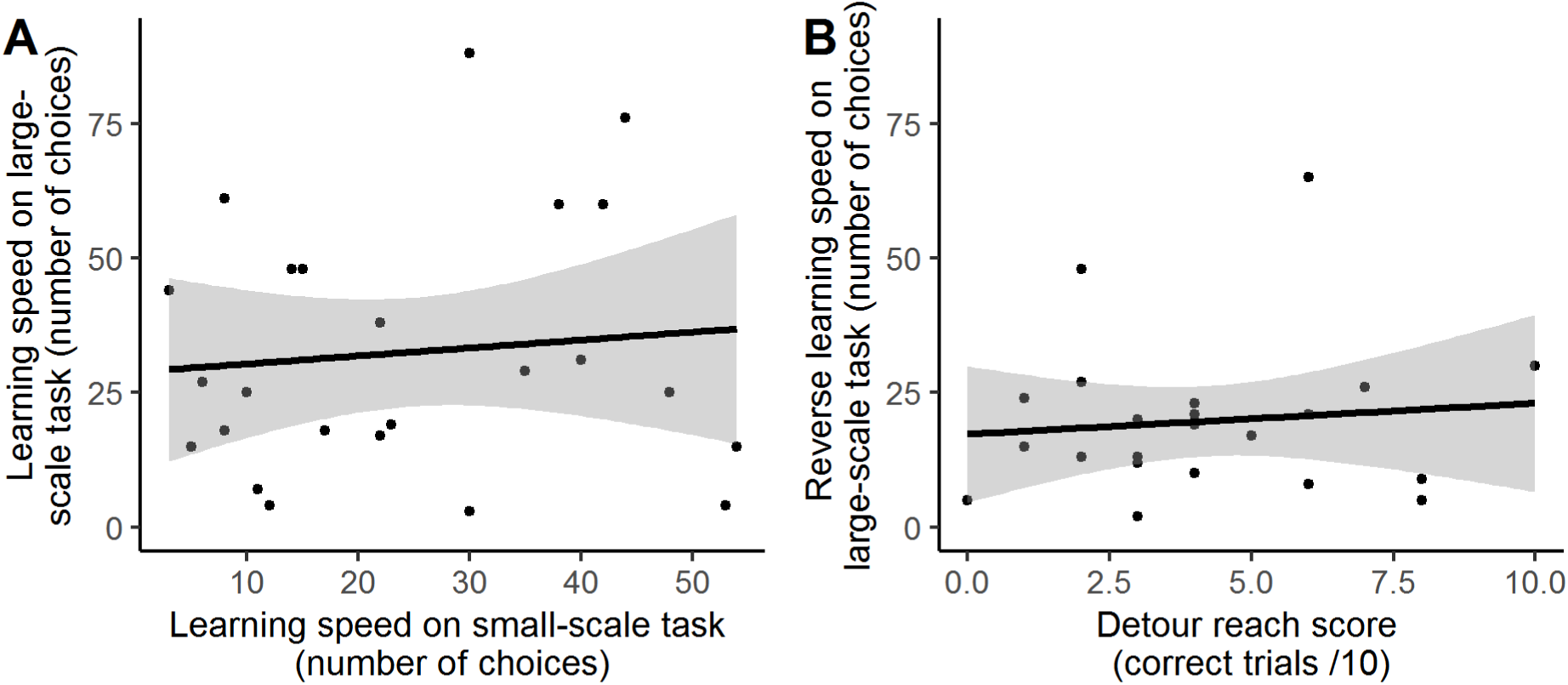
Relationship between A) performance on small-scale and large-scale spatial learning tasks, B) performance on detour reach and reversal learning tasks. The line is a fitted linear regression line and the shaded grey area is the 95% confidence interval.

On average, birds made 19.68 (SE = 3.07) choices before reverse learning (N=22), and made 3.81 (SE = 0.41) correct choices in the detour reach task (N=32). There was no evidence of a correlation between the detour reach and reversal learning (z = 0.23; tau = 0.04; p = 0.819, n=22; Figure 1.B).

## Discussion

### Spatial learning speed

We did not find a correlation between spatial learning performance in the small scale setting of the home cage and spatial learning in the larger scale setting of the experimental room. As far as we are aware, this is the first attempt to directly compare performance on a spatial learning task at different scales in non-human animals (but see Hegarty et al., 2006; Montello, 1993 for work in humans). Theoretically both tasks we used should measure spatial learning because they required individuals to use and remember specific location cues with food rewards, allowing them to return to that location more often than one would expect by chance (Olton, 1977). One possible explanation for the lack of correlation between cognitive tasks is that non-cognitive factors - for example motivation, personality, stress, and motor skills - influenced performance differently across tasks. Although we had no a priori reason to expect this might be the case, it must remain a possibility. Another plausible explanation is that different kinds of cues or strategies were used to recognise locations in our two tasks and that this confounded any correlation in cognitive performance (Morgan et al., 2014). Work in captivity has shown that several species are more reliant on cues from the geometry of the room when they have to navigate in small enclosures, and absolute direction to landmarks when they have to navigate large enclosures (Chiandetti et al., 2007; Sovrano et al., 2005, 2006). Visual information might also change at a faster pace in the home cage than in the exploration room, creating a potential bias against relying on optical-flow in the home cage. If these different perceptual abilities are either not correlated among individuals (Healy et al., 2009; Jones & Healy, 2006; Pike et al., 2018; Sovrano et al., 2003) or are independent of other processes involved with spatial learning, for example memory (Tello-Ramos et al., 2018), this could readily explain the lack of a correlation in our data (Rowe & Healy, 2014b). In this case categorising the sensory information available to the individuals could explain individuals’ spatial learning performances (Pritchard et al., 2017). Our results are in contrasts to findings that, despite having some different mechanisms, learning at different spatial scale was still found to be, to some extent, partly underlined by a common process for encoding, maintaining and transforming spatial representation (Hegarty et al., 2006; Montello, 1993).

Whatever the reason for the lack of correlation, subtle differences in experimental design may preclude meaningful comparison across studies (Thornton & Lukas, 2012). Furthermore, using only one task is unlikely to capture the expression of spatial learning in different contexts (Pritchard et al., 2017). For example, a current focus of research in food caching species is to measure spatial learning under standardised conditions, usually on a small scale, to infer performance in cache retrieval at a larger spatial scale, or to predict fitness, but it remains unclear whether observed associations relate to navigation through the environment or cache retrieval at a fine spatial scale (Healy, 2019; Healy et al., 2005, 2009; Krebs et al., 1990; McGregor & Healy, 1999). Similarly, work on parasitic cowbirds in which the females, but not males, need to locate and remember potential host nests, have found that females out-perform males in some spatial learning tasks, but not others, further suggesting that individual differences in spatial ability may depend on task design and scale of spatial location (Sherry & Guigueno, 2019). This limitation mirrors the more general problem in evolutionary ecological studies of cognition where there is often a lack of a clear link between standardised cognitive tests (e.g. problem solving) and functional behaviour (innovative foraging) under natural conditions (Morand-Ferron et al., 2016; Rowe & Healy, 2014a, 2014b; Thornton et al., 2014).

### Behavioural flexibility

We did not find any evidence of a relationship between the performance of the birds on a detour-reach and reversal learning task, despite the prediction that both measures of behavioural flexibility involve inhibitory control. This lack of correlation is in keeping with previous work in wild and captive birds, which have also used single reversals (song sparrow, *Melospiza melodia*: Boogert et al., 2011; New Zealand robin, *Petroica longipes*: Shaw et al., 2015). Our study increases the generality of this finding because these previous studies focussed on colour discrimination reversal rather than the spatial discrimination reversal we used here. There are multiple possible explanations for why these putative measures of inhibitory control may not necessarily correlate. One is that these different tasks in reality measure different components of inhibitory control (namely stopping, delaying, or withholding motor responses) (Bari & Robbins, 2013; Bray et al., 2014; Brucks et al., 2017; van Horik, Langley, Whiteside, Laker, Beardsworth, et al., 2018; Vernouillet et al., 2018). Moreover, although our results suggest that components of inhibitory control do not fall under the same general domain umbrella of behavioural flexibility, the lack of correlation may also be explained by the reversal learning task requiring a spatial learning component (Boogert et al., 2018; Brucks et al., 2017; Miyake et al., 2000). Discrepancies could also be explained by differences in motivation, which may have played a bigger role in the detour reach task as the food was visible, compared to the reversal learning task, though we note that this is likely primarily an issue if rank order differences in motivation between individuals varies across tasks. Finally, although we acknowledge that the detour-reach task has been recently criticised as a measure of inhibitory control and is subject to various non-cognitive influences, including perception, stress, prior experience (Kabadayi et al., 2018; van Horik, Langley, Whiteside, Laker, Beardsworth, et al., 2018), this is likely true of all cognitive traits (Morand-Ferron & Quinn, 2015).

## Conclusions

Our results across both task comparisons highlight that caution needs to be taken when making conclusions about learning speed or behavioural flexibility based on a single test, and that performance is highly sensitive to the context and type of task. On the one hand, if the lack of evidence for correlations reflects true cognitive differences related to either spatial learning or behavioural flexibility, then this would point towards greater complexity in the cognitive processes that drive animal navigation and behavioural plasticity. On the other hand, if the lack of correlation arises because of confounding effects which were not controlled for, or because one or both of the tasks within each domain does not measure learning or behavioural flexibility as expected, then this would point towards a common issue with experimental design used in cognitive tests. Either way, the context in which we measure cognition is therefore essential to consider, if we want to better understand causes and consequences of individual variation in cognition. Further investigation into the neurobiology related to performance in tasks which *a priori* measure the same cognitive processes may facilitate progress in validating cognitive tasks (van Horik, Langley, Whiteside, Laker, Beardsworth, et al., 2018), and distinguishing which of our interpretations are most valid. However, this approach may not often be feasible in non-model, wild animals. Pinning down the meaningful measures of individual differences in cognitive mechanisms remains a major challenge. Nevertheless, studies that aim to validate tasks, as we aimed to do here, are a step forward in understanding causes and consequences of individual variation in cognition.

## Acknowledgements

We would like to thank Sam Bayley, Ciara Sexton and Karen Cogan for capturing and releasing birds. We are also grateful to Allen Whitaker for help in coordinating the aviary building and Karen Cogan for the administrative support throughout the project.

## Supplementary Information

### All birds

In the main text, the data description and analysis exclude birds that learn on their first choice, as we cannot say whether they learned or made a choice by chance (large-scale spatial learning task n = 1; large-scale spatial reversal learning task n = 3; small-scale spatial learning task n =2).

Here we report the same analysis on all birds, without exclusion:

On average, birds made 29.04 (SE = 4.44) choices before learning the large-scale spatial learning task (N=28), and 24.37 (SE = 3.11) choices before learning the small-scale spatial learning task (N=30). Although birds needed to make more choices to learn the large-scale spatial learning task (model estimate = 0.29; C.I. = 0.18-0.39; p-value <0.001), they took slightly less time (seconds) to learn it (model estimate = −0.04; C.I. =−0.04; −0.03; p-value <0.001), compared to the small-scale spatial learning task. Birds took on average 129.06 (SE = 12.20) minutes to learn the large-scale spatial learning task, while it took them 134.57 (SE = 19.11) minutes to learn the small-scale spatial learning task. There was no evidence of a correlation between the two measures (z = 0.99; tau = 0.14; p = 0.320, n=26).

On average, birds made 17.44 (SE = 2.97) choices before reverse learning (N=25), and made 3.81 (SE = 0.41) correct choices in the detour reach task (N=32). There was no evidence of a correlation between the detour reach and reversal learning (z = 0.59; tau = 0.00; p = 0.554, n=25).

### Control birds

A similar analysis was run on the control birds, and found similar results to the correlations with all the birds included.

On average, birds assigned to the control condition made 22.11 (SE = 6.49) choices before learning the large-scale spatial learning task (N=9), and 22.44 (SE = 6.69) choices before learning the small-scale spatial learning task (N=9). There was no evidence of a correlation between the two measures (z = −1.82; tau = −0.59; p = 0.068, n=7).

On average, birds assigned to the control condition made 18.37 (SE = 5.62) choices before reverse learning (N=8), and made 3.90 (SE = 0.74) correct choices in the detour reach task (N=10). There was no evidence of a correlation between the detour reach and reversal learning (z = −0.15; tau = −0.05; p = 0.878, n=7).

## Notes

### Competing Interest Statement

The authors have declared no competing interest.

## References

Anderson, R. C., Searcy, W. A., Peters, S., Hughes, M., DuBois, A. L., & Nowicki, S. (2017). Song learning and cognitive ability are not consistently related in a songbird. Animal Cognition, 20(2), 309–320. https://doi.org/10.1007/s10071-016-1053-7

Aplin, L. M., Farine, D. R., Morand-Ferron, J., Cockburn, A., Thornton, A., & Sheldon, B. C. (2015). Experimentally induced innovations lead to persistent culture via conformity in wild birds. Nature, 518(7540), 538–541. https://doi.org/10.1038/nature13998

Ashton, B. J., Ridley, A. R., Edwards, E. K., & Thornton, A. (2018). Cognitive performance is linked to group size and affects fitness in Australian magpies. Nature, 554(7692), 364–367. https://doi.org/10.1038/nature25503

Audet, J.-N., & Lefebvre, L. (2017). What’s flexible in behavioral flexibility? Behavioral Ecology, 28(4), 943–947. https://doi.org/10.1093/beheco/arx007

Bari, A., & Robbins, T. W. (2013). Inhibition and impulsivity: Behavioral and neural basis of response control. Progress in Neurobiology, 108, 44–79. https://doi.org/10.1016/j.pneurobio.2013.06.005

Bates, D., Machler, M., Bolker, B. M., & Walker, S. C. (2015). Fitting linear mixed-effects models using lme4.

Boogert, N. J., Anderson, R. C., Peters, S., Searcy, W. A., & Nowicki, S. (2011). Song repertoire size in male song sparrows correlates with detour reaching, but not with other cognitive measures. Animal Behaviour, 81(6), 1209–1216. https://doi.org/10.1016/j.anbehav.2011.03.004

Boogert, N. J., Madden, J. R., Morand-Ferron, J., & Thornton, A. (2018). Measuring and understanding individual differences in cognition. Philosophical Transactions of the Royal Society B: Biological Sciences, 373(1756), 20170280. https://doi.org/10.1098/rstb.2017.0280

Branch, C. L., Pitera, A. M., Kozlovsky, D. Y., Bridge, E. S., & Pravosudov, V. V. (2019). Smart is the new sexy: Female mountain chickadees increase reproductive investment when mates to males with better spatial cognition. Ecology Letters, 22(6), 897–903.

Bray, E. E., MacLean, E. L., & Hare, B. A. (2014). Context specificity of inhibitory control in dogs. Animal Cognition, 17(1), 15–31. https://doi.org/10.1007/s10071-013-0633-z

Brown, V. J., & Tait, D. S. (2014). Behavioral flexibility: Attentional shifting, rule switching, and response reversal. In I. P. Stolerman & L. H. Price (Eds.), Encyclopedia of Psychopharmacology (pp. 1–7). Springer Berlin Heidelberg. https://doi.org/10.1007/978-3-642-27772-6_340-2

Brucks, D., Marshall-Pescini, S., Wallis, L. J., Huber, L., & Range, F. (2017). Measures of dogs’ inhibitory control abilities do not correlate across tasks. Frontiers in Psychology, 8, 849. https://doi.org/10.3389/fpsyg.2017.00849

Cauchard, L., Boogert, N. J., Lefebvre, L., Dubois, F., & Doligez, B. (2013). Problem-solving performance is correlated with reproductive success in a wild bird population. Animal Behaviour, 85(1), 19–26. https://doi.org/10.1016/j.anbehav.2012.10.005

Cauchoix, M., Chow, P. K. Y., van Horik, J. O., Atance, C. M., Barbeau, E. J., Barragan-Jason, G., Bize, P., Boussard, A., Buechel, S. D., Cabirol, A., Cauchard, L., Claidière, N., Dalesman, S., Devaud, J. M., Didic, M., Doligez, B., Fagot, J., Fichtel, C., Henke-von der Malsburg, J., … Morand-Ferron, J. (2018). The repeatability of cognitive performance: A meta-analysis. Philosophical Transactions of the Royal Society B: Biological Sciences, 373(1756), 20170281. https://doi.org/10.1098/rstb.2017.0281

Chiandetti, C., Regolin, L., Sovrano, V. A., & Vallortigara, G. (2007). Spatial reorientation: The effects of space size on the encoding of landmark and geometry information. Animal Cognition, 10(2), 159–168. https://doi.org/10.1007/s10071-006-0054-3

Cole, E. F., Cram, D. L., & Quinn, J. L. (2011). Individual variation in spontaneous problem-solving performance among wild great tits. Animal Behaviour, 81(2), 491–498. https://doi.org/10.1016/j.anbehav.2010.11.025

Cole, E. F., Morand-Ferron, J., Hinks, A. E., & Quinn, J. L. (2012). Cognitive ability influences reproductive life history variation in the wild. Current Biology, 22(19), 1808–1812. https://doi.org/10.1016/j.cub.2012.07.051

Cole, E. F., & Quinn, J. L. (2012). Personality and problem-solving performance explain competitive ability in the wild. Proceedings of the Royal Society B: Biological Sciences, 279(1731), 1168–1175. https://doi.org/10.1098/rspb.2011.1539

Cooke, A. C., Davidson, G. L., Reichert, M. S., & Quinn, J. L. (In prep). Non-lethal effects of predators on prey in the context of spatial learning and cognitive flexibility.

Croston, R., Branch, C. L., Kozlovsky, D. Y., Dukas, R., & Pravosudov, V. V. (2015). Heritability and the evolution of cognitive traits: Table 1. Behavioral Ecology, 26(6), 1447–1459. https://doi.org/10.1093/beheco/arv088

Dutour, M., Suzuki, T. N., & Wheatcroft, D. (2020). Great tit responses to the calls of an unfamiliar species suggest conserved perception of call ordering. Behavioral Ecology and Sociobiology, 74(3), 37. https://doi.org/10.1007/s00265-020-2820-7

Guenther, A., & Brust, V. (2017). Individual consistency in multiple cognitive performance: Behavioural versus cognitive syndromes. Animal Behaviour, 130, 119–131. https://doi.org/10.1016/j.anbehav.2017.06.011

Hartig, F. (2020). DHARMa: Residual diagnostics for hierarchical (Multi-Level / Mixed) (0.2.7) [Computer software]. https://CRAN.R-project.org/package=DHARMa

Healy, S. D. (2019). Animal cognition. Integrative Zoology, 14(2), 128–131. https://doi.org/10.1111/1749-4877.12366

Healy, S. D., Bacon, I. E., Haggis, O., Harris, A. P., & Kelley, L. A. (2009). Explanations for variation in cognitive ability: Behavioural ecology meets comparative cognition. Behavioural Processes, 80(3), 288–294. https://doi.org/10.1016/j.beproc.2008.10.002

Healy, S. D., Dekort, S., & Clayton, N. (2005). The hippocampus, spatial memory and food hoarding: A puzzle revisited. Trends in Ecology & Evolution, 20(1), 17–22. https://doi.org/10.1016/j.tree.2004.10.006

Healy, S. D., & Hurly, T. A. (1998). Rufous hummingbirds’ (*Selasphorus rufus*) memory for flowers: Patterns or actual spatial locations? Journal of Experimental Psychology, 24(4), 396–404.

Healy, S. D., & Hurly, T. A. (2004). Spatial learning and memory in birds. Brain, Behavior and Evolution, 63(4), 211–220. https://doi.org/10.1159/000076782

Hegarty, M., Montello, D. R., Richardson, A. E., Ishikawa, T., & Lovelace, K. (2006). Spatial abilities at different scales: Individual differences in aptitude-test performance and spatial-layout learning. Intelligence, 34(2), 151–176. https://doi.org/10.1016/j.intell.2005.09.005

Ihalainen, E., Lindström, L., & Mappes, J. (2007). Investigating Müllerian mimicry: Predator learning and variation in prey defences. Journal of Evolutionary Biology, 20(2), 780–791. https://doi.org/10.1111/j.1420-9101.2006.01234.x

Isden, J., Panayi, C., Dingle, C., & Madden, J. (2013). Performance in cognitive and problem-solving tasks in male spotted bowerbirds does not correlate with mating success. Animal Behaviour, 86(4), 829–838. https://doi.org/10.1016/j.anbehav.2013.07.024

Jones, C. M., & Healy, S. D. (2006). Differences in cue use and spatial memory in men and women. Proceedings of the Royal Society B: Biological Sciences, 273(1598), 2241–2247. https://doi.org/10.1098/rspb.2006.3572

Kabadayi, C., Bobrowicz, K., & Osvath, M. (2018). The detour paradigm in animal cognition. Animal Cognition, 21(1), 21–35. https://doi.org/10.1007/s10071-017-1152-0

Keagy, J., Savard, J.-F., & Borgia, G. (2009). Male satin bowerbird problem-solving ability predicts mating success. Animal Behaviour, 9.

Krebs, J. R., Healy, S. D., & Shettleworth, S. J. (1990). Spatial memory of paridae: Comparison of a storing and a non-storing species, the coal tit, Parus ater, and the great tit, P. major. Animal Behaviour, 39(6), 1127–1137. https://doi.org/10.1016/S0003-3472(05)80785-7

Langley, E. J. G., Adams, G., Beardsworth, C. E., Dawson, D. A., Laker, P. R., van Horik, J. O., Whiteside, M. A., Wilson, A. J., & Madden, J. R. (2020). Heritability and correlations among learning and inhibitory control traits. Behavioral Ecology, araa029. https://doi.org/10.1093/beheco/araa029

Loukola, O. J., Adamik, P., Adriaensen, F., Barba, E., Doligez, B., Flensted-Jensen, E., Eeva, T., Kivelä, S. M., Laaksonen, T., Morosinotto, C., Mänd, R., Niemelä, P. T., Remeš, V., Samplonius, J. M., Sebastiano, M., Senar, J. C., Slagsvold, T., Sorace, A., Tschirren, B., … Forsman, J. T. (2020). The roles of temperature, nest predators and information parasites for geographical variation in egg covering behaviour of tits (Paridae). Journal of Biogeography, jbi.13830. https://doi.org/10.1111/jbi.13830

MacLean, E. L., Hare, B., Nunn, C. L., Addessi, E., Amici, F., Anderson, R. C., Aureli, F., Baker, J. M., Bania, A. E., Barnard, A. M., Boogert, N. J., Brannon, E. M., Bray, E. E., Bray, J., Brent, L. J. N., Burkart, J. M., Call, J., Cantlon, J. F., Cheke, L. G., … Zhao, Y. (2014). The evolution of self-control. Proceedings of the National Academy of Sciences, 111(20), E2140–E2148. https://doi.org/10.1073/pnas.1323533111

McGregor, A., & Healy, S. D. (1999). Spatial accuracy in food-storing and nonstoring birds. Animal Behaviour, 58(4), 727–734. https://doi.org/10.1006/anbe.1999.1190

Miyake, A., Friedman, N. P., Emerson, M. J., Witzki, A. H., Howerter, A., & Wager, T. D. (2000). The unity and diversity ofexecutive functions and their contributions to complex “frontal lobe” tasks: A latent variable analysis. Cognitive Psychology, 41(1), 49–100. https://doi.org/10.1006/cogp.1999.0734

Montello, D. R. (1993). Scale and multiple psychologies of space. In A. U. Frank & I. Campari (Eds.), Spatial information theory: A theoretical basis for GIS (Springer, Vol. 716). https://doi.org/10.1007/3-540-57207-4_21

Morand-Ferron, J., Cole, E. F., & Quinn, J. L. (2016). Studying the evolutionary ecology of cognition in the wild: A review of practical and conceptual challenges: Evolutionary ecology of cognition in the wild. Biological Reviews, 91(2), 367–389. https://doi.org/10.1111/brv.12174

Morand-Ferron, J., Cole, E. F., Rawles, J. E. C., & Quinn, J. L. (2011). Who are the innovators? A field experiment with 2 passerine species. Behavioral Ecology, 22(6), 1241–1248. https://doi.org/10.1093/beheco/arr120

Morand-Ferron, J., & Quinn, J. L. (2015). The evolution of cognition in natural populations. Trends in Cognitive Sciences, 19(5), 235–237. https://doi.org/10.1016/j.tics.2015.03.005

Morgan, K. V., Hurly, T. A., & Healy, S. D. (2014). Individual differences in decision making by foraging hummingbirds. Behavioural Processes, 109, 195–200. https://doi.org/10.1016/j.beproc.2014.08.015

Olton, D. S. (1977). Spatial Memory. Scientific American, 236(6), 82–99.

O’Shea, W., O’Halloran, J., & Quinn, J. L. (2018). Breeding phenology, provisioning behaviour, and unusual patterns of life history variation across an anthropogenic heterogeneous landscape. Oecologia, 188(4), 953–964. https://doi.org/10.1007/s00442-018-4155-x

Pike, T. W., Ramsey, M., & Wilkinson, A. (2018). Environmentally induced changes to brain morphology predict cognitive performance. Philosophical Transactions of the Royal Society B: Biological Sciences, 373(1756), 20170287. https://doi.org/10.1098/rstb.2017.0287

Pritchard, D. J., & Healy, S. D. (2018). Taking an insect-inspired approach to bird navigation. Learning & Behavior, 46(1), 7–22. https://doi.org/10.3758/s13420-018-0314-5

Pritchard, D. J., Hurly, T. A., & Healy, S. D. (2018). Wild hummingbirds require a consistent view of landmarks to pinpoint a goal location. Animal Behaviour, 137, 83–94. https://doi.org/10.1016/j.anbehav.2018.01.014

Pritchard, D. J., Tello Ramos, M. C., Muth, F., & Healy, S. D. (2017). Treating hummingbirds as feathered bees: A case of ethological cross-pollination. Biology Letters, 13(12), 20170610. https://doi.org/10.1098/rsbl.2017.0610

Quinn, J. L., Charmantier, A., Garant, D., & Sheldon, B. C. (2006). Data depth, data completeness, and their influence on quantitative genetic estimation in two contrasting bird populations. Journal of Evolutionary Biology, 19(3), 994–1002. https://doi.org/10.1111/j.1420-9101.2006.01081.x

Quinn, J. L., Cole, E. F., Reed, T. E., & Morand-Ferron, J. (2016). Environmental and genetic determinants of innovativeness in a natural population of birds. Philosophical Transactions of the Royal Society B: Biological Sciences, 371(1690), 20150184. https://doi.org/10.1098/rstb.2015.0184

R Core Team. (2019). R: A Language and Environment for Statistical Computing (3.6.1) [Computer software]. R Foundation for Statistical Computing.

Raine, N. E., & Chittka, L. (2008). The correlation of learning speed and natural foraging success in bumble-bees. Proceedings of the Royal Society B: Biological Sciences, 275(1636), 803–808. https://doi.org/10.1098/rspb.2007.1652

Rowe, C., & Healy, S. D. (2014a). Measuring cognition will be difficult but worth it: A response to comments on Rowe and Healy. Behavioral Ecology, 25(6), 1298–1298. https://doi.org/10.1093/beheco/aru190

Rowe, C., & Healy, S. D. (2014b). Measuring variation in cognition. Behavioral Ecology, 25(6), 1287–1292. https://doi.org/10.1093/beheco/aru090

Sewall, K. B., Soha, J. A., Peters, S., & Nowicki, S. (2013). Potential trade-off between vocal ornamentation and spatial ability in a songbird. Biology Letters, 9(4), 20130344. https://doi.org/10.1098/rsbl.2013.0344

Shaw, R. C., Boogert, N. J., Clayton, N. S., & Burns, K. C. (2015). Wild psychometrics: Evidence for ‘general’ cognitive performance in wild New Zealand robins, *Petroica longipes*. Animal Behaviour, 109, 101–111. https://doi.org/10.1016/j.anbehav.2015.08.001

Shaw, R. C., & Schmelz, M. (2017). Cognitive test batteries in animal cognition research: Evaluating the past, present and future of comparative psychometrics. Animal Cognition, 20(6), 1003–1018. https://doi.org/10.1007/s10071-017-1135-1

Sherry, D. F., & Guigueno, M. F. (2019). Cognition and the brain of brood parasitic cowbirds. Integrative Zoology, 14(2), 145–157. https://doi.org/10.1111/1749-4877.12312

Soha, J. A., Peters, S., Anderson, R. C., Searcy, W. A., & Nowicki, S. (2019). Performance on tests of cognitive ability is not repeatable across years in a songbird. Animal Behaviour, 158, 281–288. https://doi.org/10.1016/j.anbehav.2019.09.020

Sonnenberg, B. R., Branch, C. L., Pitera, A. M., Bridge, E., & Pravosudov, V. V. (2019). Natural Selection and Spatial Cognition in Wild Food-Caching Mountain Chickadees. Current Biology, 29(4), 670-676.e3. https://doi.org/10.1016/j.cub.2019.01.006

Sovrano, V. A., Bisazza, A., & Vallortigara, G. (2003). Modularity as a fish (Xenotoca eiseni) views it: Conjoining geometric and nongeometric information for spatial reorientation. Journal of Experimental Psychology: Animal Behavior Processes, 29(3), 199–210. https://doi.org/10.1037/0097-7403.29.3.199

Sovrano, V. A., Bisazza, A., & Vallortigara, G. (2005). Animals’ use of landmarks and metric information to reorient: Effects of the size of the experimental space. Cognition, 97(2), 121–133. https://doi.org/10.1016/j.cognition.2004.08.003

Sovrano, V. A., Bisazza, A., & Vallortigara, G. (2006). How fish do geometry in large and in small spaces. Animal Cognition, 10(1), 47–54. https://doi.org/10.1007/s10071-006-0029-4

Tello-Ramos, M. C., Branch, C. L., Pitera, A. M., Kozlovsky, D. Y., Bridge, E. S., & Pravosudov, V. V. (2018). Memory in wild mountain chickadees from different elevations: Comparing first-year birds with older survivors. Animal Behaviour, 137, 149–160. https://doi.org/10.1016/j.anbehav.2017.12.019

Thornton, A., Isden, J., & Madden, J. R. (2014). Toward wild psychometrics: Linking individual cognitive differences to fitness. Behavioral Ecology, 25(6), 1299–1301. https://doi.org/10.1093/beheco/aru095

Thornton, A., & Lukas, D. (2012). Individual variation in cognitive performance: Developmental and evolutionary perspectives. Philosophical Transactions of the Royal Society B: Biological Sciences, 367(1603), 2773–2783. https://doi.org/10.1098/rstb.2012.0214

van Horik, J. O., Langley, E. J. G., Whiteside, M. A., Laker, P. R., Beardsworth, C. E., & Madden, J. R. (2018). Do detour tasks provide accurate assays of inhibitory control? Proceedings of the Royal Society B: Biological Sciences, 285(1875), 20180150. https://doi.org/10.1098/rspb.2018.0150

van Horik, J. O., Langley, E. J. G., Whiteside, M. A., Laker, P. R., & Madden, J. R. (2018). Intra-individual variation in performance on novel variants of similar tasks influences single factor explanations of general cognitive processes. Royal Society Open Science, 5(7), 171919. https://doi.org/10.1098/rsos.171919

van Horik, J. O., Langley, E. J. G., Whiteside, M. A., & Madden, J. R. (2019). A single factor explanation for associative learning performance on colour discrimination problems in common pheasants (Phasianus colchicus). Intelligence, 74, 53–61. https://doi.org/10.1016/j.intell.2018.07.001

Vernouillet, A. A. A., Stiles, L. R., Andrew McCausland, J., & Kelly, D. M. (2018). Individual performance across motoric self-regulation tasks are not correlated for pet dogs. Learning & Behavior, 46(4), 522–536. https://doi.org/10.3758/s13420-018-0354-x

Völter, C. J., Tinklenberg, B., Call, J., & Seed, A. M. (2018). Comparative psychometrics: Establishing what differs is central to understanding what evolves. Philosophical Transactions of the Royal Society B: Biological Sciences, 373(1756), 20170283. https://doi.org/10.1098/rstb.2017.0283

